# Detection and Whole Genome of Japanese Encephalitis Virus Genotype III strains isolated from Indian Pig and Mosquito vector

**DOI:** 10.1101/2022.11.03.515130

**Authors:** Seema R. Pegu, Pranab J. Das, Joyshikh Sonowal, Gyanendra S. Sengar, Rajib Deb, Swaraj Rajkhowa, Manjisa Choudhury, B. R. Gulati, V. K. Gupta

## Abstract

Japanese encephalitis virus(JEV) are globally prevalent as deadly pathogens in human and animals including pig, horse and cattle. Japanese encephalitis (JE) still remains as an important cause of epidemic encephalitis worldwide and exists in a zoonotic transmission cycle. Assam is one of the highly endemic state for JE in India. In the present study, to understand the epidemiological status of JE circulating in pigs and mosquito particularly in Assam, India, molecular detection of JEV and the complete genome sequencing of the JEV isolates from pig and mosquito were done. The complete genome analysis of two JEV isolates from pig and mosquito were revealed7 and 21 numbers of unique points polymorphism of nucleotide during alignment of the sequences with other available sequences, respectively. Phylogenetic analysis revealed that the isolates of present investigation were belongs to genotype III and closely related with the strains of neighbouring country China. This study represents the transboundary nature of the JEV genotype III circulation and maintained the same genotype through mosquito-swine transmission cycles.

## Introduction

Japanese encephalitis (JE) is a common mosquito born flaviviral encephalitis in Southeast Asian countries. JE virus (JEV) belongs to the family *Flaviviridae* and is principally a disease occurring in rural agricultural areas maintained in an enzootic cycle in involving water birds, domestic pigs and zoophilic Culex mosquito (Schuh et al., 2013). Human and horses are the dead end host, in which JEV causes acute encephalitis with the onset of acute febrile illness and alteration in mental status such as confusion, inability to talk, disorientation, seizures and coma(Singh et al., 2020). Hence, JE is a cause of public health concern (Morita et al., 2015; Singh et al., 2020). Pigs and water birds are the amplifying host of the JEV without any clinical signs, except abortion and still birth in pregnant sows and gilts (Morita et al., 2015). The incidence of Japanese encephalitis in recent times has shown increasing trend in India and has become a major public health problem (Bandyopadhyay et al., 2013; Singh et al., 2020). Assam is the north-eastern (NE) state of India having largest pig population in the country with abundance of rainfall and rice cultivated land which makes Assam the most vulnerable state for JE incidences (Datey et al., 2020; Medhi et al., 2017; Singh et al., 2020). In Assam, JEV was first reported in the year 1976 from Lakhimpur district and since then incidences of JE have markedly increased in the recent years (Hazarika, 1991; Phukan et al., 2004). Furthermore, the presence of JEV infected pigs increases the chances of spill over of infection to human, especially when the mosquito density is high.

Previous studies indicated that JEV can be classified into five genotype (I to V) based on the diversity of the nucleotide sequence of E protein gene with most isolates classified as genotype GI or genotype GIII (Zheng et al., 2012). Although numerous encephalitis etiology studies have confirmed the importance of this virus, little information on the genotype circulating strain has been reported from this part of India in amplifying host of JE. Resurgence of JE cases in human population in India during last few years highlights the need for JEV surveillance in the amplifying host. In this study, we report the two isolation and full genome sequencing of two strains of JE virus from pig and mosquito vector and make a comparison with recent isolates of JE virus from pig and mosquito.

## Materials & Methods

### Collection of samples

Selection of study area and collection of pig samples were based on a pilot survey and retrospective data on the availability of pig population/pig farms/slaughter points in Kamrup and Jorhat district of Assam, the area of study was identified (Fig. 1). A total of 112 nos. of pig samples (tissue, cerebrospinal fluid& blood) were collected from 8 pig farms and 4 slaughter points of specified districts of Assam from 2018 to 2020. A suitable questionnaire was prepared for collecting baseline information from the pig farms/slaughter points regarding age, breed, parity, no. of piglet born, any abortion of stillbirth cases, any other disease condition in the farm etc.

**Fig. 1:**
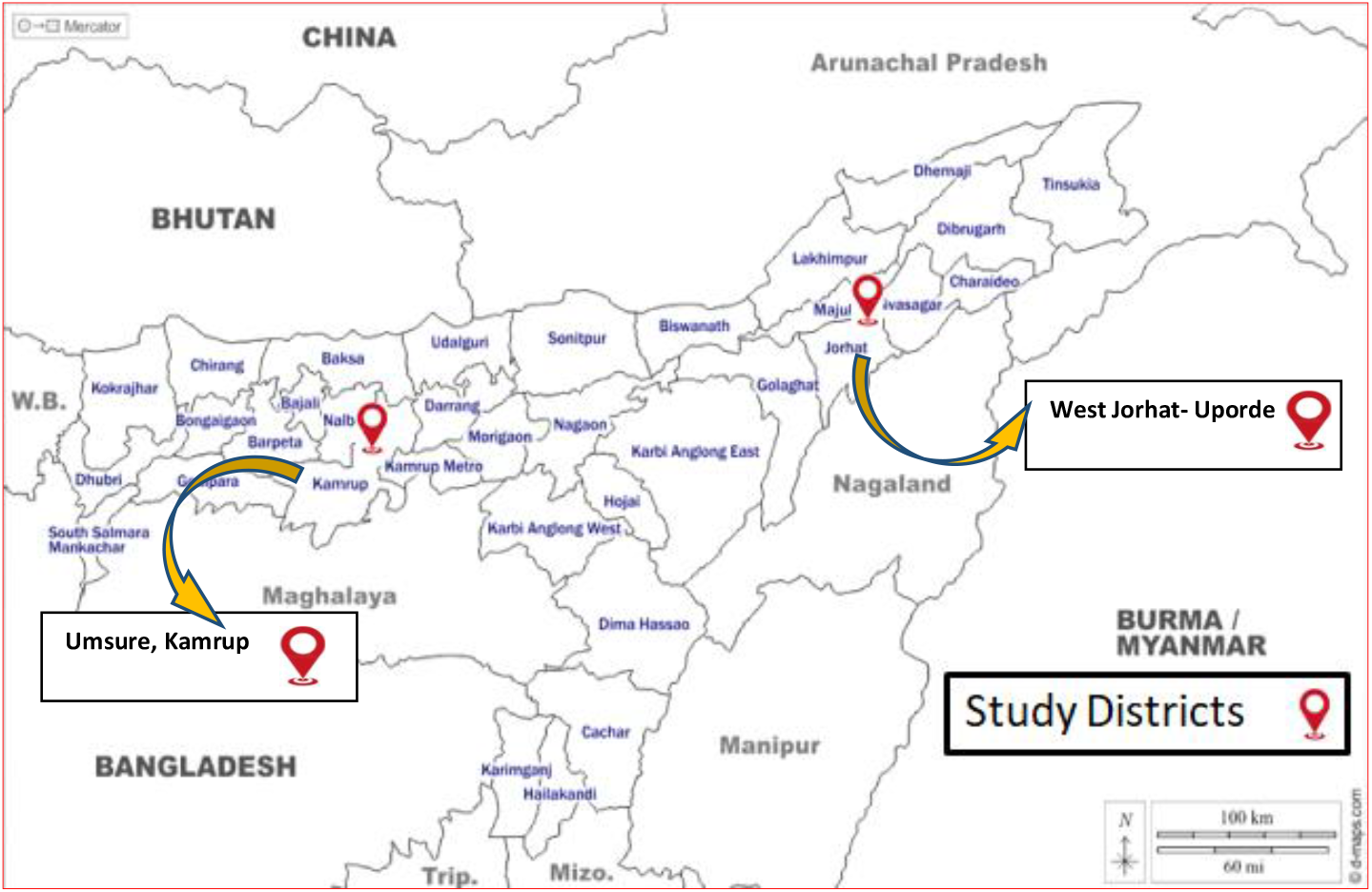
Demonstration of the geographical area where samples or specimens were collected for the current study. Here, the Assam, India, districts of Kamrup and Jorhat were chosen due to their high sales of pigs and pig products and location of JE hotspot.

Simultaneously, a total of 8124 mosquitoes were collected from Kamrup and Jorhat districts in accordance with the areas of pig farms during the dusk period of the day. Mosquitoes were collected by aspirator, sweeping net and with the help of CDC trap light inside and outside the pig farm. The collected female mosquitoes were sorted, preserved in separate plastic vials and identified under stereoscopic binocular microscope following taxonomic key.

### Isolation of viral RNA, synthesis of cDNA and amplicon generation

Viral RNA of JEV was extracted from blood and tissue samples collected from pigs and mosquito samples (Mosquitoes were pooled as approx. 30 mosquitoes/pool) using QIAamp Viral RNA Mini Kit (QIAGEN, Hilden, Germany) as per manufacturer’s instruction. RNA was converted into complementary DNA (cDNA) using a RevertAid First Strand cDNA Synthesis Kit (Cat no: K1622, Thermo Scientific™, Lithuania - European Union) with random hexamer primers. The synthesized cDNA was used to amplify the partial envelope protein gene of JEV @ 236bp by PCR using the forward primer 5’-TTACTCAGCGCAAGTAGGAGCGTCTCAAG-3’ and reverse primer 5’-ATGCCGTGCTTGAGGGGGACG-3’ (Yeh et al., 2010). A specific real time RT-PCR assay was used for the detection of JEV gene. Cycling conditions comprised Initial denaturation 95 °C for 10 s, 40 cycles of 95 ° for 15 s and 55 °C for 30 s and Amplification curves were visualised in AriaMx v1.7 software. All the samples detected by real time PCR was subjected to conventional PCR as described previously. Conventional PCR amplification was performed in thermal cycler under the following conditions: cycle at 95 ° for 2 min (initial denaturation), 40 cycles (denaturation at 95 ° for 30 s, annealing at 65 ° for 30 s, and extension at 72 ° for 35 s), and cycle of final extension at 72 ° for 10 min. Five microliters of amplified PCR products were separated by 2 % ethidium bromide strained agarose gel electrophoresis at 100 V for 25 min. 100 bp plus DNA Marker (Cat no.: SM1153, Thermo Scientific) was used as standard and the amplified products were visualized using ultraviolet light transilluminator.

### Next-generation sequencing (NGS)/ Whole-genome sequencing

The whole genome sequencing was done from the two positive samples from pig and one positive sample from mosquito pool. Whole genome sequencing of the positive viral RNA samples were outsourced (Genotypic Technology Pvt. Ltd, India). Library preparation and sequencing were prepared with Illumina-compatible NEBNext® Ultra™ II using Directional RNA Library Prep Kit (New England BioLabs, MA, USA). Viral total RNA was taken for fragmentation and priming. Fragmented and primed RNA was further subjected to first strand synthesis followed by second strand synthesis. The double stranded cDNA was purified using JetSeq Beads (Bioline, Cat # BIO-68031). Purified cDNA was end-repaired, adenylated and ligated to Illumina multiplex barcode adapters (Universal adapter: 5’-AATGATACGGCGACCACCGAGATCTACACTCTTTCCCTACACGACGCTCTTCCGA TCT-3’ and Index adapter: 5’-GATCGGAAGAGCACACGTCTGAACTCCAGTCAC [INDEX] ATCTCGTATGCCGTCTTCTGCTTG-3’) as per NEBNext® Ultra™ II Directional RNA Library Prep protocol followed by second strand excision using USER enzyme at 37 °C for 15mins. Adapter ligated cDNA was purified using JetSeq Beads and was subjected to 11 cycles for Indexing-(98°C for 30sec, cycling (98°C for 10sec, 65°C for 75 sec) and 65°C for 5mins) to enrich the adapter-ligated fragments. Final PCR product (sequencing libraries) was purified with JetSeq Beads, followed by library quality control check. Illumina-compatible sequencing libraries were quantified by Qubit Fluorometer (Thermo Fisher Scientific, MA, USA) and its fragment size distribution was analyzed on Agilent 2200 Tape Station.

### Genomic alignment, Single nucleotide polymorphism and phylogenetic Analysis

JEV complete genome sequences were obtained from the NCBI database and the accession no. of all the sequences were presented in supplementary table 1. The mosquito and pig isolates were analysed independently in two groups. Our samples were compared to other fully sequenced JEV strains in the database using the neighbour-joining (NJ) method and the ClustalW multiple alignment tool in MegAlign software. The aligned sequences underwent SNP identification and genetic relationship determination with our mosquito and pig isolates, separately. Also, we have isolated the common SNPs present in JEV isolate from pig of India, separately. To establish the genetic relationship of JEV isolates from pig and mosquito, we constructed phylogenetic tree based on Whole gene sequences.

## Results

### 3.1 Laboratory testing of JE case specimens

#### 3.1.1 Gross and histopathological findings

In our study investigated and found that a sow in her first farrowing gave birth to four stillborn piglets at full term (Fig. 2A) and tested positive for JEV. The vector mosquito collected from the pig sheds and nearby drainage and waterlogged areas revealed the prevalence of *Culex tritaeniorhynchus* mosquitoes in abundance during the evening and early morning period (Fig. 2B). On the post-mortem examination of all the stillborn piglets showed varying degrees of congestion in the lung, liver, heart, spleen and lymph nodes (Fig. 2C). Upon opening the head, two piglets showed encephalitis in the cerebrum (Fig. 2D). In histopathological investigation, the most significant microscopic lesions found in the brain were neuronophagi, satellitosis and vacuolation in the cerebrum (Fig. 2E), as well as lymphoid depletion in the peripheral lymph nodes (Fig. 2F).

**Fig. 2:**
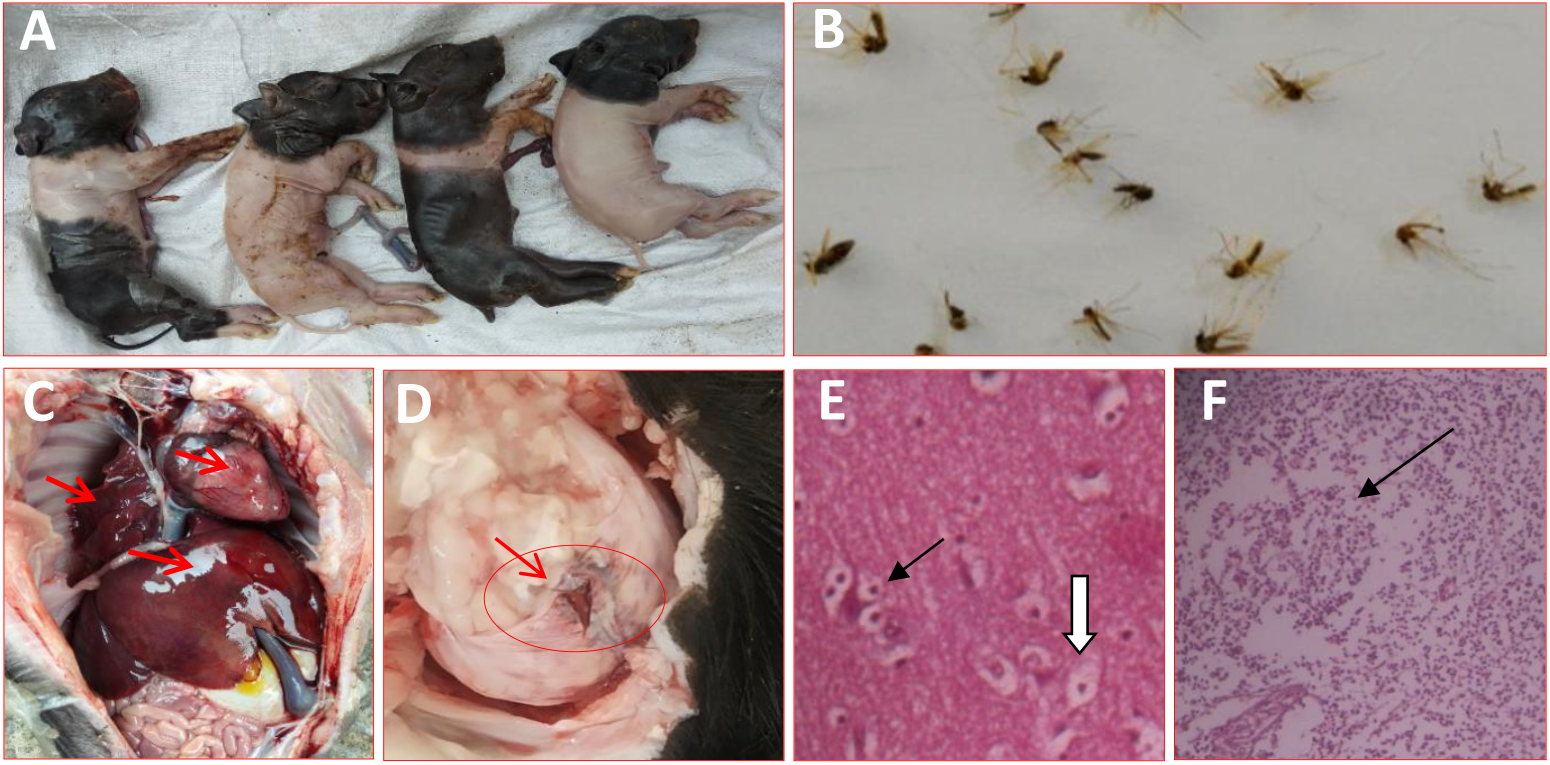
Japanese encephalitis (JE) case specimens of the current study. Here, A. Stillborn fetuses in a sow due to JE infection during pregnancy, B. A pool of *Culex tritaeniorhynchus* mosquito, Major vector of JEV transmission, C. Congestion in the internal organs (arrow), D. Encephalitis in the brain in a stillborn foetus affected with JE (arrow), E. Neuropathology in the cerebrum of a JEV-infected stillborn piglet. Neuronophagia and satellites (arrow), neuronal vacuolation (arrowheads) H and E, 20× magnification. F. Lymphoid depletion in the lymphoid follicle of lymph node (arrow) H and E, 20× magnification.

#### 3.1.2 Molecular diagnosis/ Viral gene detection (RT-PCR and real time RT-PCR)

Out of 112 pig samples, JEV was positive in 11 samples (Table 1). A total of 271 pools of mosquitoes were constituted by the end of collection based of species and sex and tested for JEV by real-time RT-PCR. Out of 271 pool two pools of *Culex tritaeniorhynchus* and *Culex gelidus*, respectively from female mosquito pools were found to be positive (Table 2). The representative screened samples were shown in Fig. 3 (Real time PCR) and Fig. 4 (Conventional PCR).

**Table 1:** Description of JEV positive samples collected from pigs from Jorhat and Kamrup District

| Sample details |  |  |  |  | Results |  | Genotype |
| --- | --- | --- | --- | --- | --- | --- | --- |
| Animal ID | Age | Sex | Type | Location | Real-time RT-PCR | Conventional RT-PCR |  |
| Kr-31 | 62d | F | Blood | Rani, Kamrup | +ve | +ve | GIII |
| Kr-51 | 75 | F | Blood | Rani, Kamrup | +ve | +ve | GIII |
| Kr-71 | 114 | F | Blood | Rani, Kamrup | +ve | +ve | GIII |
| Kr-8 | Foetus | F | Tissue | Rani, Kamrup | +ve | +ve | GIII |
| Ksd-17 | Foetus | F | Tissue | Sundubi, Kamrup | +ve | +ve | GIII |
| Ksd-29 | Foetus | F | Tissue | Sundubi, Kamrup | +ve | +ve | GIII |
| Kg-9 | 8M | M | Tissue | Gorchuk, Kamrup | +ve | +ve | GIII |
| Kl-6 | 1YR | M | Tissue | Lakhra, Kamrup | +ve | +ve | GIII |
| Jnm-46 | 9m | m | Tissue | Namdeuri, West Jorhat | +ve | +ve | GIII |
| Jnm-102 | 12m | F | Tissue | Namduri, West Jorhat | +ve | +ve | GIII |
| Jup-98 | 8m | F | Tissue | Upper deuri, West Jorhat | +ve | +ve | GIII |
| Jup-11 | 8m | F | Tissue | Upper deuri, West Jorhat | +ve | +ve | GIII |

**Fig. 3:**
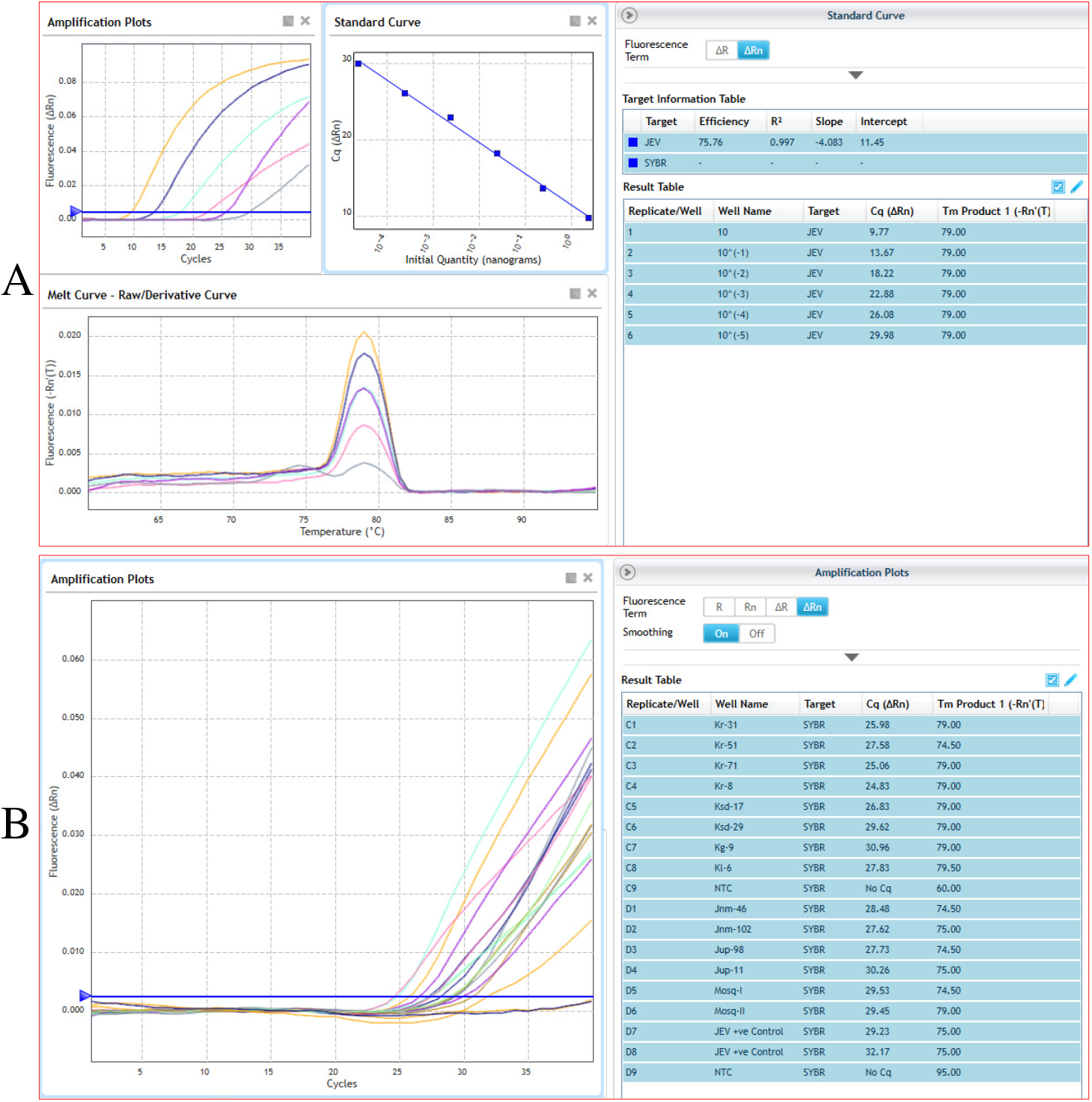
Real time PCR for detection of Japanese encephalitis virus (JEV) from different pig and mosquito samples. Here, A. shows the standard curve analysis of a representative JEV positive control sample, where 10-fold dilution factor of the sample represents 75.76% of efficiency upto 6^th^ dilution (10^−5^). The 6^th^ and 7^th^ dilution of samples were used as representative positive control for detection of JEV concentration in the present study; B. shows the presence of JEV in representative test pig and mosquito samples. These indicated that the positive samples were contains JEV cDNA/ RNA in each respective reaction.

**Fig. 4:**
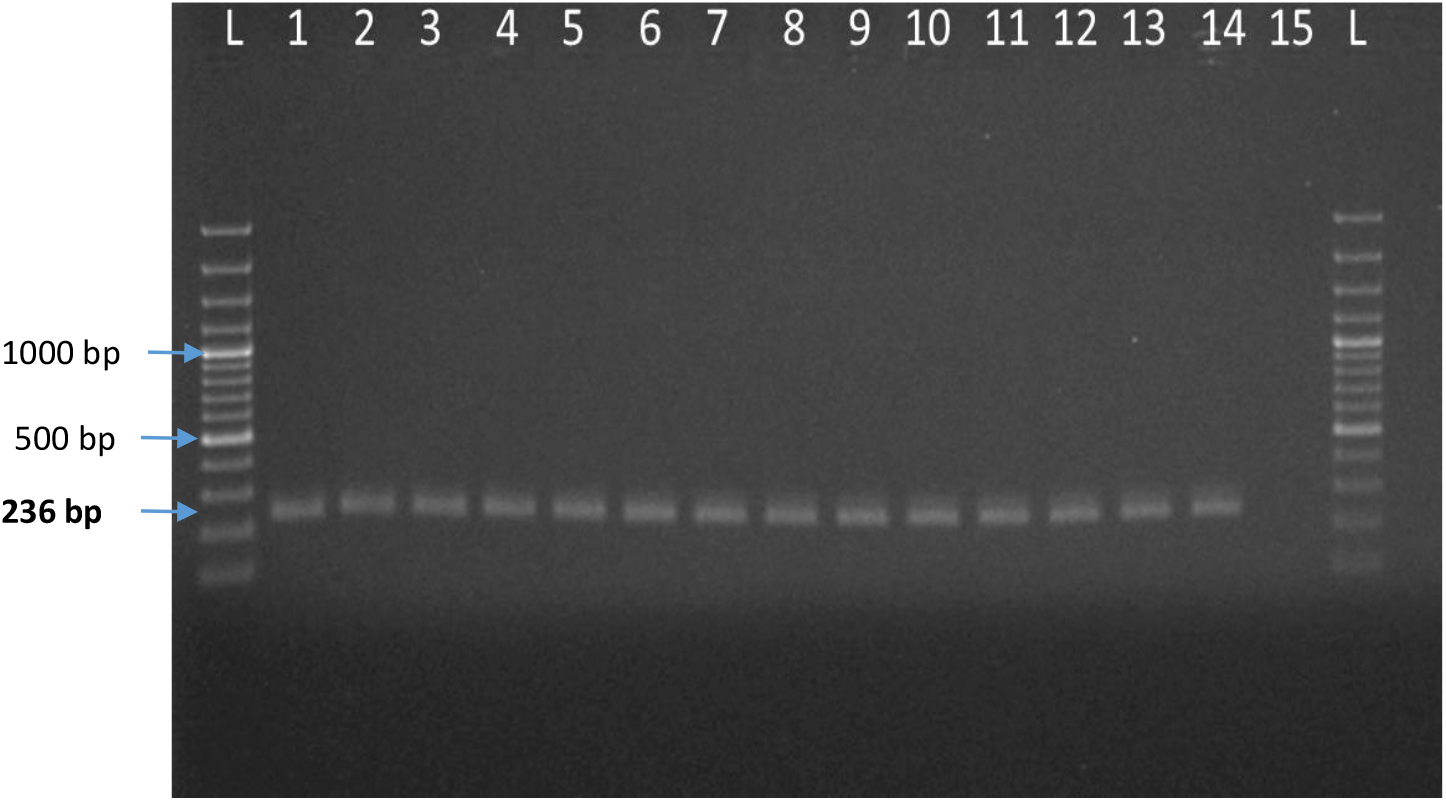
Agarose gel electrophoresis of JEV partial E gene amplified products isolated from pig and mosquito. Here, L: 100bp plus ladder, 1: Positive control of JEV, 2 to 12: Test samples of JEV isolated from pig sources, 13 and 14: Test samples of JEV isolated from mosquito, 15: Negative control

**Fig.5:**
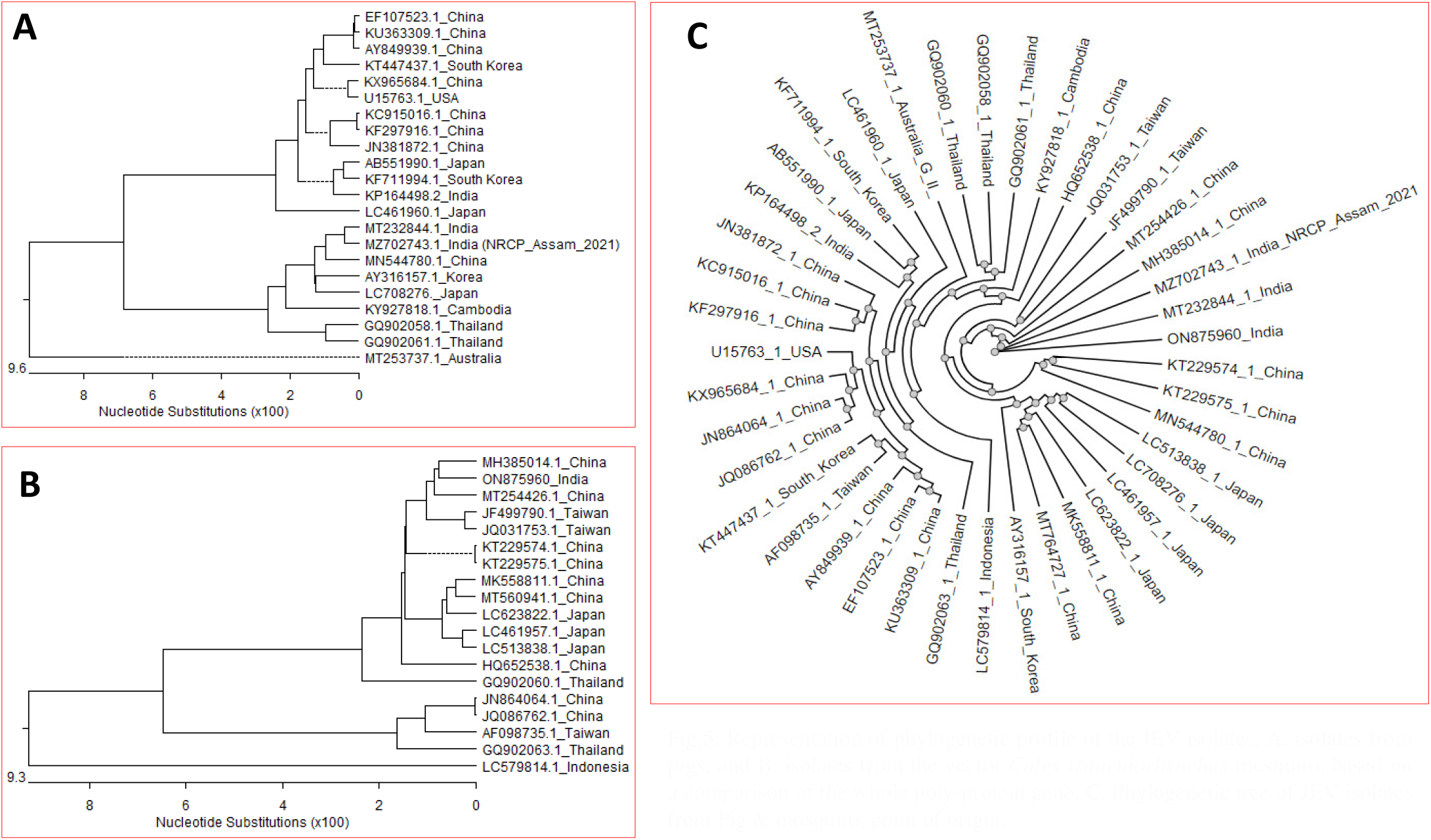
Representation of phylogenetic profile of the JEV isolates, A. isolates from pigs, and B. isolates from the vector *Culex tritaeniorhynchus* mosquito, based on a comparison of the whole poly-protein gene, C. Phylogenetic tree of JEV isolates from Pig & mosquito, point of origin.

### 3.2 Whole genome sequencing

The complete genome of the JEV detected in a pig sample and in mosquito pool sample from Assam (JEV/Assam/Pig), India was sequenced. The raw sequence data of the viral whole genome isolated from pig tissue sample was assembled and annotated using bioinformatics software and was found to be 10,966 nucleotides(nt) long with 97 nt at the 5′ untranslated region (UTR), as well as a 10,299-nt open reading frame (single) corresponding to 3,432 amino acids excluding stop codon and 570 nt of the 3′ UTR.

The positive mosquito pool samples were also sequenced and analysed to be found to be 10,965 nucleotides(nt) long with 96 nt at the 5′ untranslated region (UTR), as well as a 10,299-nt open reading frame (single) corresponding to 3,432 amino acids excluding stop codon and 570 nt of the 3′ UTR.

### 3.3 Phylogenetic analysis

The sequence alignments and phylogenetic analysis revealed that the virus belongs to genotype III. The NCBI BLAST analysis revealed that the JEV/Assam/Pig genome sequence shared highest similarity (99.2%)with Indian JEV strain isolated from pig (MT232844) followed by 98.3% of China variant (MN544780) (Table 2). The complete genome of the mosquito vector showed a high nucleotide homology revealed (98%) to reference strain 057434 (EF623988). The NCBI Blast mosquito sequences revealed 98.5% (MH385014:MT254426) of China. Phylogenetic analysis of both amplifying host and vector showing region specific similarity with the vector and host (Fig.3). Both the sequences of JEV identified in this study is also showing same trend having the greatest similarity of mosquito sequence (ON875960 and MZ782743). The radial phylogenetic analysis of all the sequences shows that. Distance matrix of all vector and host under this study Assam mosquito shows highest similarity with 99.4% with china pig and mosquito and Assam pig showed highest similarity with Indian pigs and Assam mosquitoes (Supplementary data).

**Table 2:**
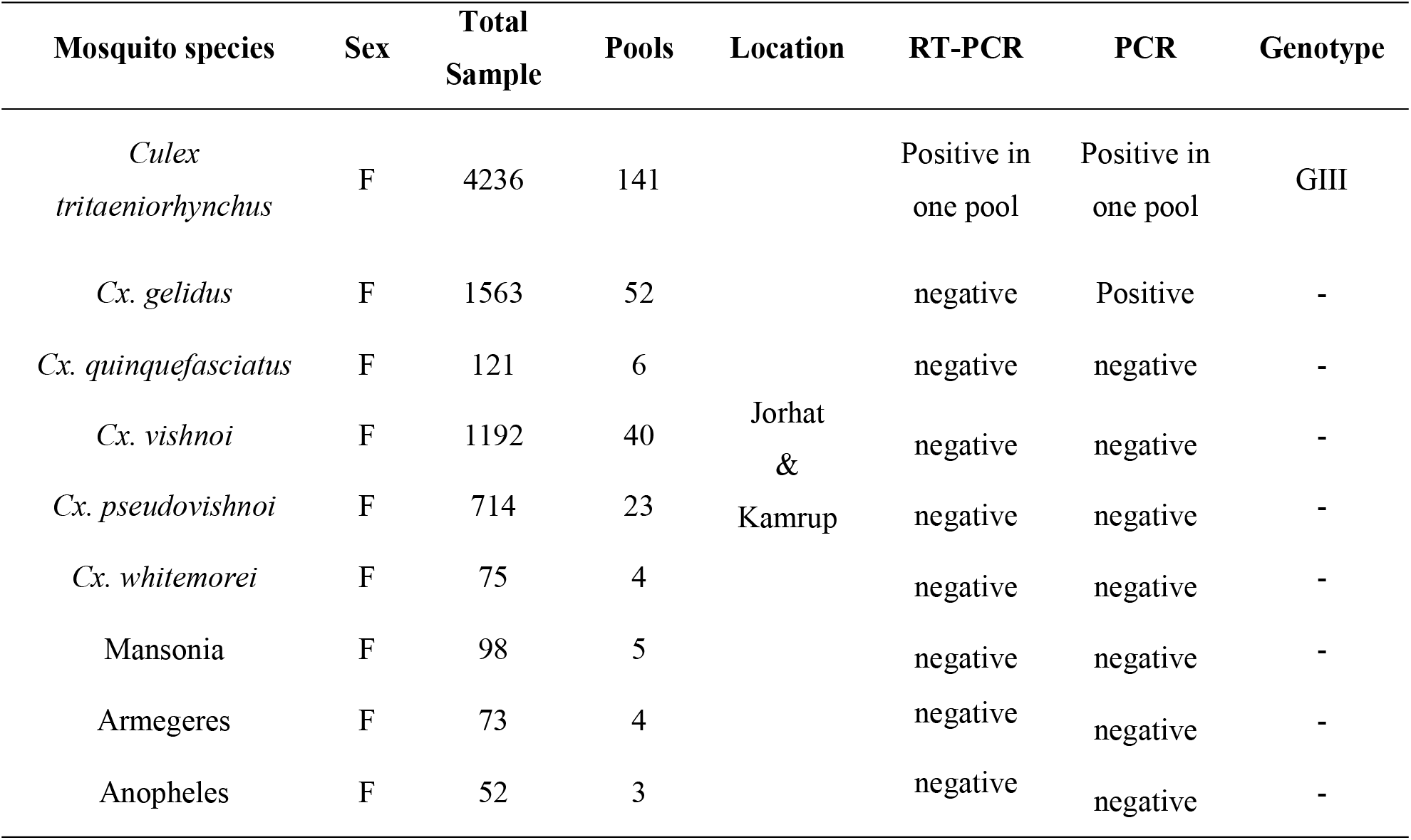
List of mosquito species identified with result of JEV screening

### 3.4 Unique polymorphism of nucleotides

A total 20 no.s of unique polymorphism of nucleotides were found in polyprotein gene of JEV isolated from mosquito (Fig. 6a, Supplementary table 2). In case of JEV isolated from pigs, 7 no.s of unique polymorphism of nucleotides were found in polyprotein gene (Fig. 6b, Supplementary table 3). Additionally, 15no.s of unique polymorphism of nucleotides were commonly observed in polyprotein gene of JEV strains isolated from pigs of India (MZ702743.1 and MT232844.1) from other reference genes (Fig. 6c, Supplementary table 4).

**Fig. 6:**
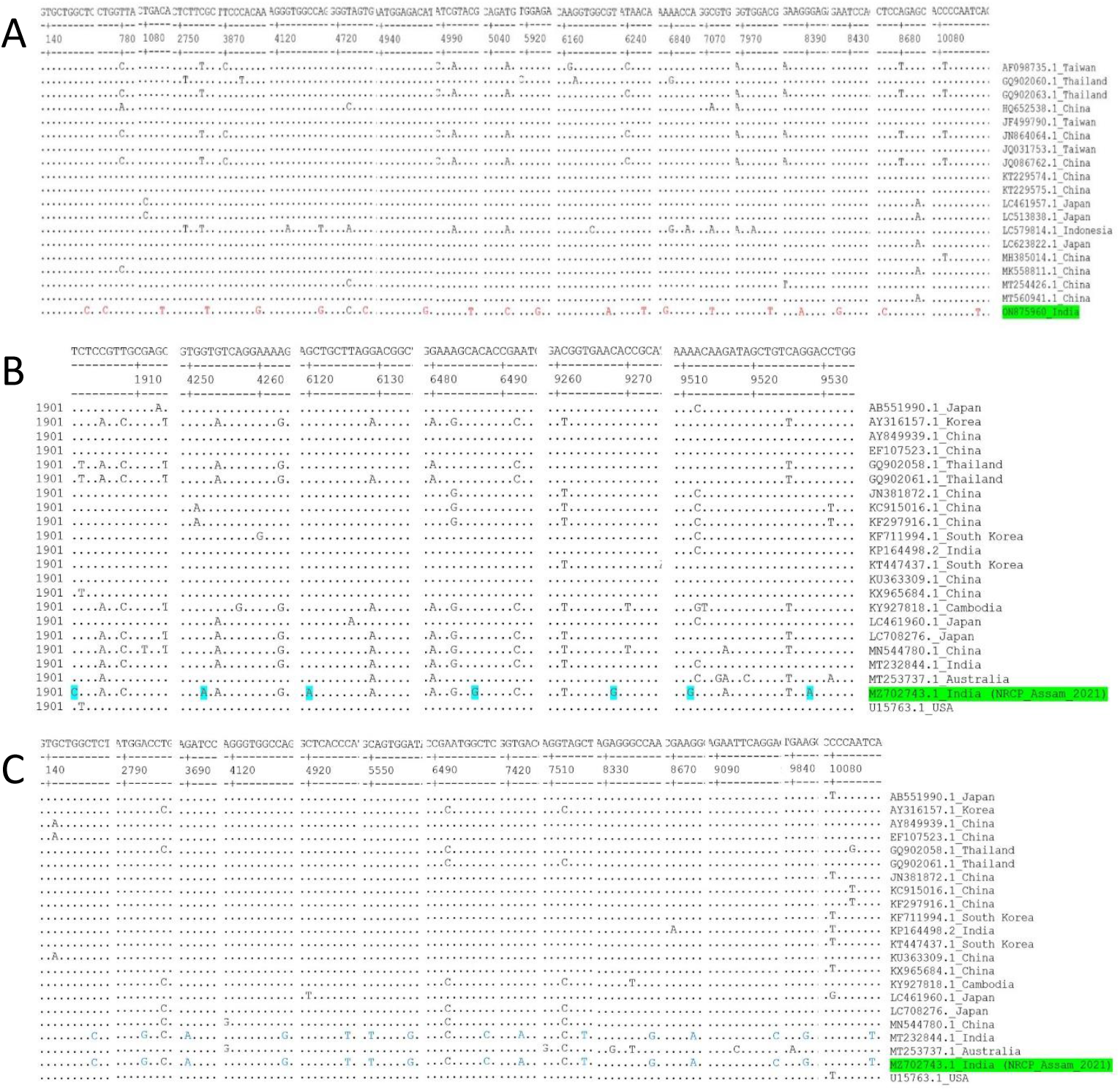
Nucleotide sequence alignment of the polyprotein genes of JEV isolates from A. mosquito, B. pig and SNP identification in Indian isolate of current study. C. Common and unique SNPs of polyprotein genes of JEV isolates from two Indian strain. Residues identical to the consensus are indicated by dots.

## Discussion

JEV is an RNA virus that have a high potential for evolution due to their lack of repair mechanisms that would otherwise act during the replication of their genome (Domingo and Holland, 1997; Holland et al., 1982). Extensive surveillance of amplifying hosts and vectors assists in understanding geographical migration and genotype shift in JEV (Liu et al., 2013). The current work represents a constrained example of JEV molecular evolution in nature. JEV is still a serious but neglected public health problem in India and is the most frequent etiology of meningeo-encephalitis in human of several Asian countries (Gong et al., 2010; Shailendra et al., 2014). In North Eastern region India, large JE epidemics occurs in human during the monsoon season when mosquito density is abundant. There has been a recent increase of JE/Acute Encephalitis Syndrome (AES) cases in Assam, which have migrated from upper Assam to every adjoining districts and there have been reports of confirmed cases and mortality throughout the state (Dev et al., 2015). In the present study, from 2018 to 2020 a total of 112 nos. of pig samples were collected, we detected JEV by RT PCR in 11 pig samples. A total of 8124 mosquitoes were collected from the Jorhat and Kamrup districts in accordance with the areas pig samples were collected. A total nine (9) mosquito species were identified, including *Culex tritaeniorhynchus, Culex vishnoi, Culex pseudovishnoi, Culex gelidus, Culex whitemoorei*, Mansonia. The most prevalent species was *Culex tritaeniorhynchus*, followed by *Cx. Gelidas* and the *Cx. Vishnui, Culex pseudovishnoi* and *Cx. quinquefasciatus* subgroup. A total of 278 pools of mosquitoes consisting 20-30 mosquitoes in one pool were constituted every fortnight of collection based on species and tested for JEV by RT-PCR. Out of all the tested samples only in one pools of *Cx. tritaeniorhynchus* captured in August 2019 from Kamrup rural was found to be positive subjected for WGS.

The polyprotein gene is the complete ORF of JEV flanked by 5′ and 3′ untranslated regions (UTRs) and the polyprotein consisting of structural proteins such as capsid [C], membrane [prM/M], and envelope [E] and non-structural proteins such as NS1, NS2A, NS2B, NS3, NS4A, NS4B, and NS5 (Ng et al., 2017). The polyprotein gene sequences of JEV stain isolated from pig and isolated from mosquito were compared separately for the genetic relationship via phylogenetic tree with other stains of respective JEV isolated from pig and mosquitoes, and showed that a number of nucleotide substitutions (unique polymorphism) that were scattered throughout the polyprotein gene. The analysis of JEV from pig revealed the circulation of JEV genotype III in the pig population of Assam, India and this is region specific. This is India’s first study to disclose the entire genomic sequences of JEV isolated from mosquitos. However, the differences in the sequence variation did not result in the changes in genotype among the strains but virulence might be varying. Thus the polyprotein gene sequence suitable for genotyping as well as monitoring gene evolution.

The JEV isolates from pigs and mosquitoes of the present study showed less mutation and variation in the point of origin analysis of different JEV sequences, indicating that our isolates were the most ancestral. The phylogenetic analysis of JEV (MZ702743.1) isolated from pig showed that they were closely related to the isolates of India (MT232844.1) followed by the isolate of neighbouring country China (MN544780.1). On the other hand, analysis indicated that JEV isolated form mosquito for the first time from India (ON875960.1) was closely related to the strains of neighbouring country China (MH385014.1 and MT254426.1). This investigation indicates the transboundary nature of the JE. Since North East states of India are located on the border of China, Myanmar, Bhutan and Bangladesh, this investigation not only provides a basis for the prevalence of JEV in this region, but also for neighbouring countries. Thus the data from this study may aid in the design of effective control strategies for each of these regions as well as it will provide future impact on development of a new therapeutics and vaccine which against JEV includes genotype III strains.

## Conclusion

In conclusion, our report clearly showed that JEV full genome sequencing from pig and mosquito vector belongs to Genotype III circulating in this region which is same with the virus circulating in the human. It is worth mentioning that *Culex tritaeniorhynchus* is the most important vector for JEV transmission in India. Evidence of JEV transmission in this region (in pigs and mosquitoes) may have potential implications for future spread of JEV in human and other animal populations. Additional survey of JEV in pig, human and mosquito vector in NE region India is necessary to better understand strategic control of the disease.

## Authors contribution

Concept and Designing of manuscript: SRP, PJD and JS. Wet-lab work: SRP, MC, GSS and RD. Dry lab work: PJD and JS. Manuscript drafting: JS, SRP and PJD. Proof reading and editing: RD, SR, BRG, VKG, SRP, PJD, JS, and GSS. All authors revised and approved the final version of manuscript.

## Data Availability Statement

The data that support this study will be shared upon reasonable request to the corresponding author.

